# Prediction of coronavirus 3C-like protease cleavage sites using machine-learning algorithms

**DOI:** 10.1101/2021.08.15.456422

**Authors:** Huiting Chen, Zhaozhong Zhu, Ye Qiu, Xingyi Ge, Heping Zheng, Yousong Peng

## Abstract

The coronavirus 3C-like (3CL) protease is a Cysteine protease. It plays an important role in viral infection and immune escape by not only cleaving the viral polyprotein ORF1ab at 11 sites, but also cleaving the host proteins. However, there is still a lack of effective tools for determining the cleavage sites of the 3CL protease. This study systematically investigated the diversity of the cleavage sites of the coronavirus 3CL protease on the viral polyprotein, and found that the cleavage motif were highly conserved for viruses in the genera of *Alphacoronavirus, Betacoronavirus* and *Gammacoronavirus*. Strong residue preferences were observed at the neighboring positions of the cleavage sites. A random forest (RF) model was built to predict the cleavage sites of the coronavirus 3CL protease based on the representation of residues at cleavage site and neighboring positions by amino acid indexes, and the model achieved an AUC of 0.96 in cross-validations. The RF model was further tested on an independent test dataset composed of cleavage sites on host proteins, and achieved an AUC of 0.88 and a prediction precision of 0.80 when considering the accessibility of the cleavage site. Then, 1,079 human proteins were predicted to be cleaved by the 3CL protease by the RF model. These proteins were enriched in pathways related to neurodegenerative diseases and pathogen infection. Finally, a user-friendly online server named 3CLP was built to predict the cleavage sites of the coronavirus 3CL protease based on the RF model. Overall, the study not only provides an effective tool for identifying the cleavage sites of the 3CL protease, but also provides insights into the molecular mechanism underlying the pathogenicity of coronaviruses.

## Introduction

The coronavirus is a kind of positive-sense single-stranded RNA viruses (Hartenian et al., 2020). It can be grouped into four genera including *Alphacoronavirus, Betacoronavirus, Gammacoronavirus*, and *Deltacoronavirus*. Seven coronaviruses have been reported to infect humans, including HCoV-NL63 and HCoV-229E (*Alphacoronavirus*), HCoV-OC43, HCoV-HKU1, severe acute respiratory syndrome coronavirus (SARS-CoV), Middle East respiratory syndrome coronavirus (MERS-CoV) and SARS-CoV-2 (*Betacoronavirus*) (Chen et al., 2020; Hartenian et al., 2020). Among them, SARS-CoV, MERS-CoV and SARS-CoV-2 are highly pathogenic and lethal (Gralinski et al., 2013; Chafekar and Fielding, 2018; Hu et al., 2021). Especially, the current pandemic caused by the SARS-CoV-2 has resulted in 200,840,180 human infections and 4,265,903 deaths globally as of August 6^th^, 2021 (WHO, 2021). How to effectively control the coronavirus is a great challenge for humans.

The coronavirus has a genome ranging from 27-32 kb in size (Cui et al., 2019). Most coronaviruses share a similar genomic structure which includes a polyprotein ORF1ab, four structural proteins (S, E, M and N), and a variable number of accessory proteins (Satija and Lal, 2007). The polyprotein ORF1ab could be cleaved into 16 non-structural proteins (NSPs) by the viral proteases which are NSP3/papain-like protease and NSP5/3C-like protease (Klemm et al., 2020). The 3C-like (3CL) protease, a typical Cysteine protease, cleaves the ORF1ab at 11 sites and produces 11 NSPs (NSP4-NSP16) (Arya et al., 2021). These individual NSPs participate in multiple critical processes of viral infection such as the viral genome replication and transcription (Snijder et al., 2016; Arya et al., 2021). Besides, the 3CL protease can also cleave the host proteins (Wang et al., 2016; Zhu et al., 2017a; Zhu et al., 2017b; Chen et al., 2019; Zhu et al., 2020). A recent study by Moustaqil et al. found that the 3CL protease of SARS-CoV-2 could directly cleave TAB1 and NLRP12, which may provide a molecular mechanism for enhanced production of cytokines and inflammatory response observed in COVID-19 patients (Moustaqil et al., 2021). Due to the important role of 3CL protease in viral infection, it has been taken as a critical target for antiviral drug development (Anand et al., 2003; Fu et al., 2020; Vuong et al., 2020).

The cleavage sites of the 3CL protease are relatively conserved. Previous studies have shown that the first position in the upstream of the cleavage site, defined as P1 according to Schechter and Berger’s study (Schechter and Berger, 1967), was highly conserved with the amino acid (AA) Q (Anand et al., 2003; Fang et al., 2010). Besides, other positions near the cleavage site also showed strong preferences to some AAs (Anand et al., 2003; Chuck et al., 2010). For example, P2, P3 and P4 preferred the high-hydrophobicity residues, positively charged residues, and small hydrophobic residues, respectively, while the downstream positions P1’ and P2’ both preferred small residues (Chuck et al., 2011). However, considering the large diversity of coronaviruses (Cui et al., 2019), there is still a lack of a systematic study towards the diversity of the cleavage sites by the coronavirus 3CL protease.

Lots of coronavirus polyproteins lack of annotations in the public databases due to a lack of effective tools for determining the cleavage sites on the polyprotein. Besides, only a few host proteins were experimentally determined to be cleaved by the viral 3CL protease. It is in great need to develop more effective methods for determining the cleavage sites of the coronavirus 3CL protease. There are currently two kinds of computational methods for determining the cleavage sites of the virus protease. The first is the machine-learning methods (Singh and Su, 2016; Stanley et al., 2020). For example, Kiemer et al. developed a neural network model NetCorona to predict the cleavage sites of the coronavirus 3CL protease with high accuracy (Kiemer et al., 2004). Unfortunately, only 77 cleavage sites from seven full-length coronavirus genomes were used to train the model, which may lead to potential bias. The other is the homology-based method which infers the cleavage sites of polyproteins by sequence alignment to the reference sequences with known cleavage sites (Larsen et al., 2020). Although it is accurate for sequences which are highly similar to reference sequences, it can be hardly applied for those with large diversification to reference sequences, and is unable to annotate the host proteins cleaved by the 3CL protease.

This work systematically investigated the diversity of the cleavage sites of 3CL protease in coronaviruses and built a random forest (RF) model for predicting the cleavage sites of the coronavirus 3CL protease with high accuracy; the RF model was further tested by the experimentally determined cleavage sites in host proteins; then, the RF model was used to predict the cleavage sites of the coronavirus 3CL protease on the human proteome; finally, a user-friendly online server named 3CLP was built to predict the cleavage sites of the coronavirus 3CL protease based on the RF model. The work would not only help understand the specificity of the coronavirus 3CL protease, but also facilitate the annotation of proteins cleaved by the coronavirus 3CL protease.

## Materials and methods

### The coronavirus cleavage sites

At least one polyprotein (ORF1ab) sequence with the known cleavage sites of the 3CL protease were obtained for 14 coronavirus species in three genera including *Alphacoronavirus, Betacoronavirus* and *Gammacoronavirus* from the NCBI RefSeq and protein databases on April 15^th^,2021 (Table S1). For each coronavirus species with the known cleavage sites on at least one polyprotein, the polyprotein sequences were obtained from the NCBI protein database and were aligned with MAFFT (version 7.427) (Katoh and Standley, 2013). Since the cleavage of polyproteins by 3CL protease is important for viral infection, the cleavage sites on polyproteins of the same viral species are hypothesized to be highly conserved and were obtained with the homology-based method (Larsen et al., 2020). A window of 20 AAs centered on each cleavage site was obtained from each sequence. Previous studies have shown that the P1 position was highly conserved with Q (Anand et al., 2003; Fang et al., 2010). Therefore, only the windows with Q in P1 were kept. The windows from viruses of the same genus were combined together and were de-duplicated. The number of unique windows in each genus was listed in Table S2.

### The data for modeling

Because positions P2-P4 (P1 was excluded because it was supposed to be completely conserved) and P1’-P2’ were more conserved than other positions (Figure 1), the motifs containing residues in these positions were further extracted and were defined as the cleavage motifs. A total of 905 cleavage motifs were obtained from the genera of *Alphacoronavirus, Betacoronavirus* and *Gammacoronavirus*. They were then de-duplicated, which resulted in 265 unique cleavage motifs. They were taken as positive samples in the modeling. To obtain the negative samples, the Qs in polyprotein sequences of 14 coronavirus species mentioned above were identified except those in the cleavage sites; then, for each Q, a non-cleavage motif containing the neighboring three AAs in the upstream of Q and two AAs in the downstream of Q was built. A total of 6,828 non-cleavage motifs were obtained. Based on the one-hot encoding, these non-cleavage motifs were grouped into 265 clusters by the k-means method using the module of sklearn.cluster in Python (version 3.7) (Pedregosa et al., 2011). One motif was randomly selected from each cluster, which led to 265 negative samples.

**Figure 1.**
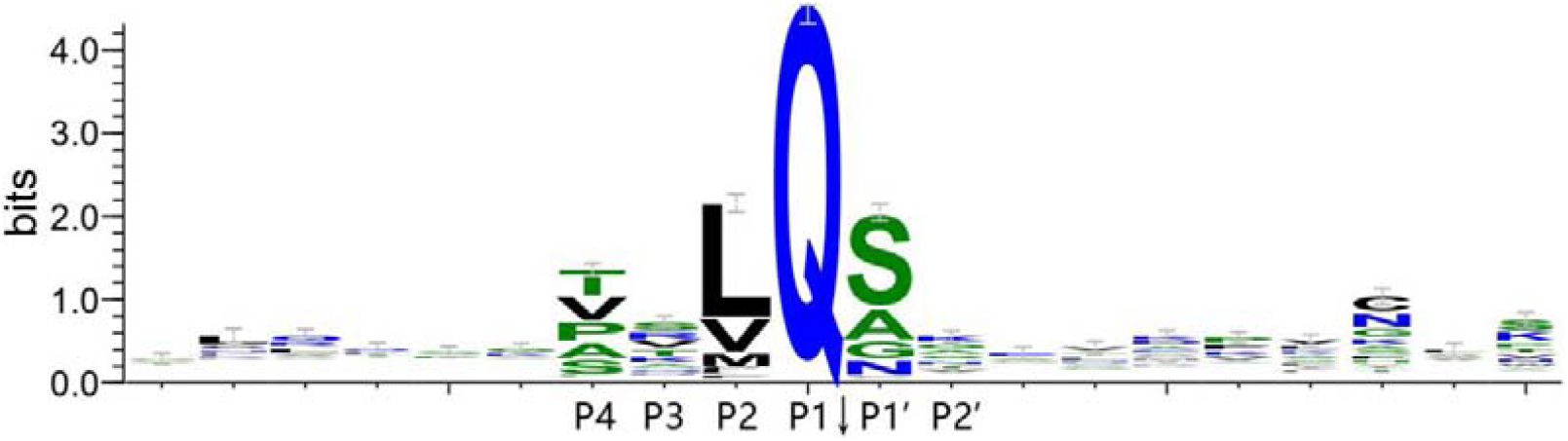
The logo for sequences around the cleavage sites of the coronavirus 3CL protease.

### The AA indexes

A total of 566 AA indexes were obtained from the AAindex database (version 9.2) on November 18^th^, 2020 (Kawashima et al., 2008).

### The human proteome and the protein structures

The human proteome was obtained from the SwissProt database in UniProt on June 29^th^, 2021. The protein sequences of 20,386 human proteins were obtained. The location of human proteins were inferred based on the subcellular location provided by the UniProt database. The protein structures were obtained from the AlphaFold Protein Structure Database on July 27^th^, 2021 (Tunyasuvunakool et al., 2021).

### Logo of sequences centered the cleavage sites

The logo of sequences centered the cleavage sites was generated with WebLogo 3 using the default parameters on April 16^th^, 2021 (Crooks et al., 2004).

### Machine-learning modeling with random forest, support vector machine and naive bayes

Three machine-learning algorithms, random forest (RF), support vector machine (SVM) and naive bayes (NB), were used to predict the cleavage sites of the 3CL protease and were achieved with functions of RandomForestClassifier, svm.SVC, GaussianNB, respectively, in the package of sklearn in Python (version 3.7) (Pedregosa et al., 2011). Five times of five-fold cross-validations were used to evaluate the performance of the machine-learning algorithms. The AUC, accuracy, sensitivity, specificity, false positive rate (FPR) and precision were used to evaluate the model performance. The AUC was calculated using the module of sklearn.metrics (Pedregosa et al., 2011); the accuracy, sensitivity, specificity, FPR and precision were calculated based on the confusion matrix as the follows:

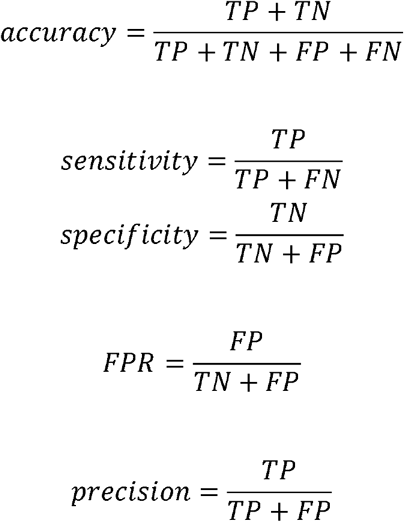

in which the TP, TN, FP and FN referred to true positive, true negative, false positive and false negative, respectively.

### The Principal Component Analysis (PCA)

The PCA of the AA indexes were achieved using the module of sklearn.decomposition in Python (version 3.7) (Pedregosa et al., 2011).

### The calculation of solvent accessible surface area (SASA) of residues in the cleavage sites

For human proteins, the residue SASA was calculated by the tool of FreeSASA (version 2.1.0) (Mitternacht, 2016) based on the protein structures; for proteins of other species, the SASA of residues in the cleavage sites was predicted by the tool of SPOT-1D-Single (version 1.0) (Singh et al., 2021).

### Functional enrichment analysis of human genes

The KEGG pathway and GO enrichment analysis was conducted with functions of enrichKEGG and enrichGO in the package clusterProfiler (version 3.18.1) in R (version 4.0.5) (Yu et al., 2012). All the KEGG pathways and GO terms with q-values less than 0.05 were considered as significant enrichment.

### Statistical analysis

All the statistical analyses in this study were conducted in R (version 4.0.5). The Wilcoxon rank-sum test was used to compare the sample means in this study and was conducted with the function of *wilcox*.*test()* in R.

## Results

### The diversity of cleavage sites of the coronavirus 3CL protease

The 3CL protease has 11 cleavage sites on the polyprotein ORF1ab of coronaviruses (Snijder et al., 2016). The cleavage sites of 3CL protease on polyproteins were obtained from 14 coronavirus species in three genera including *Alphacoronavirus, Betacoronavirus* and *Gammacoronavirus*. The logos of sequences around the cleavage sites for three genera (Figure 1 and Figure S1) showed similar residue conservation levels and residue preferences. Besides the P1, the P2, P1’ and P4 were the most conserved sites for all three genera. On the position of P2, the AAs of L, M and V were most conserved; on the P1’, the AAs of S, A, G and N were most conserved; on the P4, the AAs of T, V, P, A and S were most conserved. When combined together, the P1-P4 and P1’-P2’ were more conserved than other positions (Figure 1). Because the P1 were supposed to be completely conserved, the motif containing the P2-P4 and P1’-P2’ (defined as the cleavage motif) was kept for further analysis (Figure 2).

**Figure 2.**
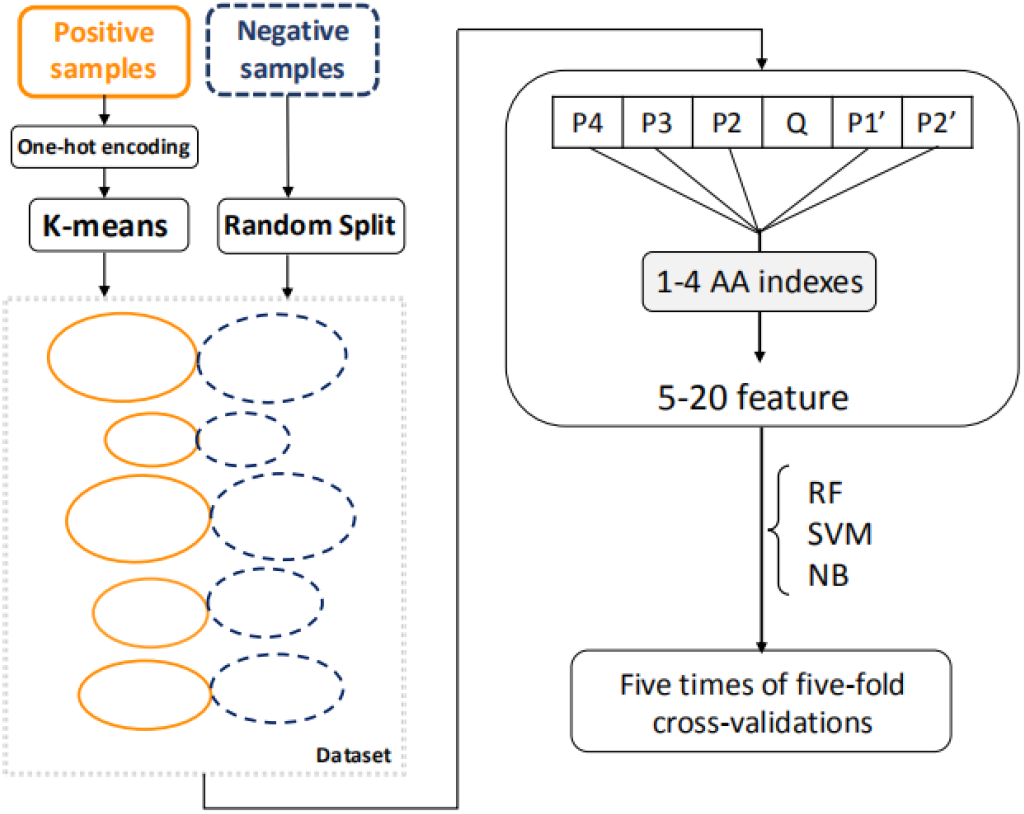
The work flow of the modeling process.

### Prediction of the cleavage sites of the 3CL protease

A total of 265 cleavage motifs (positive samples) and equal number of non-cleavage motifs (negative samples, see Materials and methods) were obtained to build the machine-learning model for predicting the cleavage sites of the coronavirus 3CL protease. The work flow of the modeling process was shown in Figure 2.

Firstly, the positive samples were encoded with the one-hot method, and were clustered into five groups by the k-means method. To ensure the balance of the positive and negative samples in the training and validation process, the negative samples were randomly separated into five groups to match the positive sample groups. The above processes were repeated five times and five datasets were generated. The size of each group in five datasets was listed in Table 1.

**Table 1.** The size of five groups in each dataset. The positive group and the corresponding negative group had the same size.

|  | Dataset1 | Dataset2 | Dataset3 | Dataset4 | Dataset5 |
| --- | --- | --- | --- | --- | --- |
|  | Pos/Neg | Pos/Neg | Pos/Neg | Pos/Neg | Pos/Neg |
| Group 1 | 99/99 | 38/38 | 48/48 | 78/78 | 58/58 |
| Group 2 | 58/58 | 77/77 | 75/75 | 54/54 | 77/77 |
| Group 3 | 30/30 | 52/52 | 39/39 | 79/79 | 79/79 |
| Group 4 | 58/58 | 41/41 | 72/72 | 34/34 | 20/20 |
| Group 5 | 20/20 | 57/57 | 31/31 | 20/20 | 31/31 |

Secondly, each AA on the positive or negative sample (the cleavage or non-cleavage motif) was encoded by one to four AA indexes, which led to five to twenty features for each sample.

Thirdly, three machine-learning algorithms including the RF, SVM and NB were used to build the model for predicting the cleavage sites of the 3CL protease, and five times of five-fold cross-validations were used to evaluate and compare the predictive performance of the algorithms. When using one AA index in the modeling, there were a total of 566 models for each algorithm. The RF models had a median AUC of 0.88, which were significantly higher than those of both the SVM and NB models (Figure 3A). Therefore, the RF algorithm was used in the further modeling.

**Figure 3.**
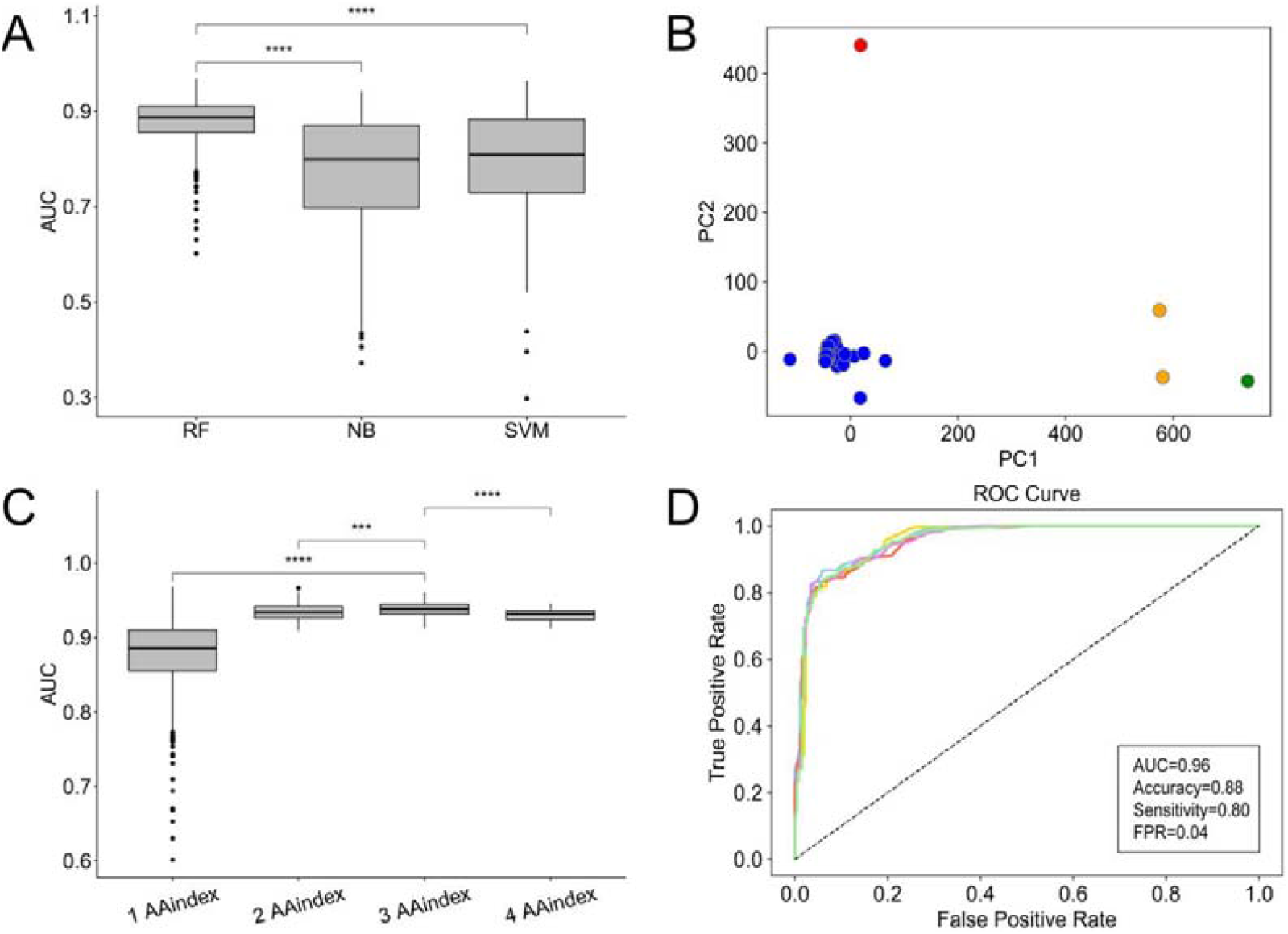
Prediction of cleavage sites of 3CL protease with the machine-learning algorithms. (A) Comparison of model performances with RF, NB and SVM algorithms when using one AA index; (B) Visualization of the first two components of the top 10% AA indexes in the PCA analysis; (C) Comparison of performances of RF models with one to four AA indexes; (D) The ROC and model performances of the best RF model using three AA indexes. ***, p-value < 0.001; ****, p-value < 0.0001.

To improve the model performance, the top 10% AA indexes (58 AA indexes) in the RF models were analyzed with the PCA method. The first and second components were visualized in Figure 3B. Four AA index clusters were obtained by the k-means clustering. To reduce the co-linearity of features in the modeling, combination of AA indexes was conducted by cluster. For example, when using two AA indexes in the modeling, two AA indexes were randomly selected from two different clusters independently. The RF models using all possible combinations of two, three and four AA indexes were built and evaluated. As shown in Figure 3C, the RF models with two AA indexes had higher AUCs than those with one AA index; the model performances were further improved when using three AA indexes; however, the model performances were decreased when using four AA indexes. Overall, the RF models using three AA indexes performed significantly better than those with one, two or four AA indexes. The RF model which performed best among all models using three AA indexes had an AUC of 0.96. More specifically, the accuracy, sensitivity, FPR of the model were 0.88, 0.80, and 0.04, respectively. The best RF model used the AA indexes of MEEJ800102, BIOV880102 and FASG760101, which referred to “the retention coefficient in high-pressure liquid chromatography”, “Information value for accessibility” and “Molecular weight”, respectively.

### Validation and application of the model

To test the RF model in prediction of the cleavage sites of the coronavirus 3CL protease, a small test dataset was manually curated from previous literatures (Table S3) (Wang et al., 2016; Zhu et al., 2017a; Zhu et al., 2017b; Chen et al., 2019; Zhu et al., 2020; Moustaqil et al., 2021). The test dataset contained 12 experimentally validated cleavage sites on seven host proteins from human, cat, pig and mouse. Except the AA of Qs in these cleavage sites, a total of 350 Qs in these proteins were hypothesized to constitute the non-cleavage sites. The RF model was tested on the test dataset and predicted 19 positive samples (taking 0.5 as the cutoff), among which 6 were experimentally validated (Table S4). The AUC, prediction accuracy, sensitivity and false positive rate were 0.88, 0.95, 0.50 and 0.04, respectively. When taking 0.95 as the cutoff for determining the predicted positive sample, only 6 positive samples were predicted, and 4 of them were experimentally validated. The false positive rate was decreased to 0.006, and the prediction precision was increased to 0.67 (Table S5). Because the cleavage sites must be accessible (Blom et al., 1996), the predicted cleavage sites could be further filtered by the solvent accessible surface area (SASA). Only the accessible cleavage sites during which the SASA of the residues in both the P1 and P1’ have greater than 10 Å^2^ were kept, which led to 5 predicted cleavage sites when taking 0.95 as the cutoff. The false positive rate was further decreased to 0.003 and the prediction precision was increased to 0.8 (Table 2).

**Table 2.**
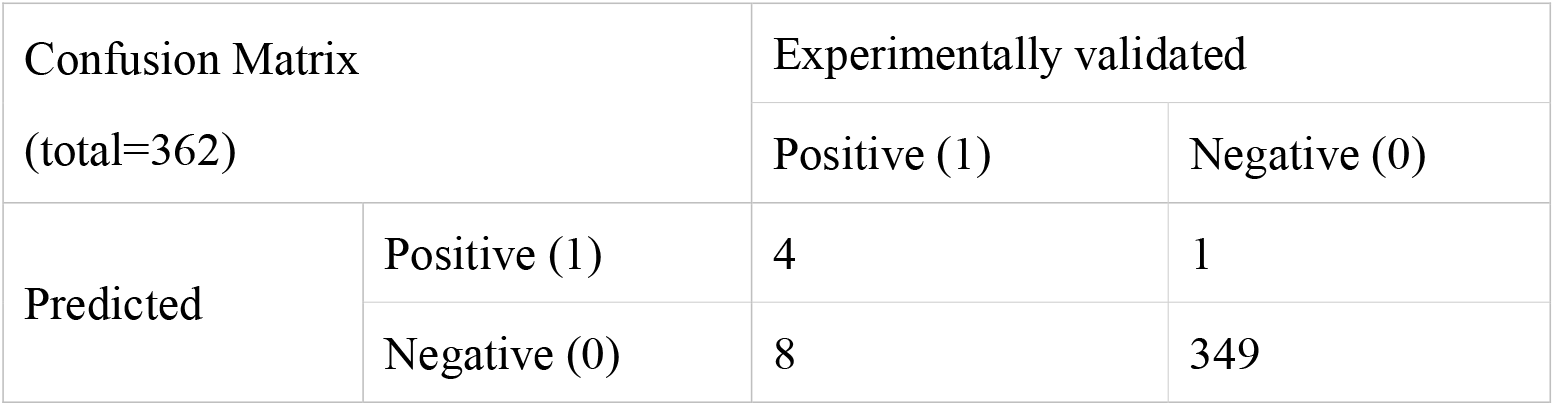
The prediction performances of the RF model on the test dataset when taking 0.95 as the cutoff and considering the accessibility of the cleavage site.

Then, the RF model was used to predict the potential cleavage sites on human proteins by the coronavirus 3CL protease. Only the proteins in the cytoplasm were considered since they are most likely to be cleaved by the coronavirus 3CL protease. To reduce the FPR and increase the precision, the cutoff for determining the positive was set to be 0.95 and only the accessible cleavage sites were considered. A total of 1,079 human proteins were predicted to be cleaved by the coronavirus 3CL protease with 1,420 cleavage sites. Most of human proteins had only one predicted cleavage sites. Some proteins had more than five cleavage sites, such as the Abnormal spindle-like microcephaly-associated protein (UniProtKB : Q8IZT6) and Baculoviral IAP repeat-containing protein 6 (UniProtKB : Q9NR09). The GO enrichment analysis of the human proteins which were predicted to be cleaved by the coronavirus 3CL protease showed that in the domain of Biological Process, they were enriched in processes of organization, assembly, movement, localization, and so on (Figure 4A and Table S6); in the domain of Cellular Component, they were enriched in microtubule, spindle, cell leading edge, and so on (Figure 4B and Table S6); in the domain of Molecular Function, they were enriched in binding, kinase activity, GTPase regulator activity, and so on (Figure 4C and Table S6). The KEGG enrichment analysis showed that these proteins were enriched in pathways related to neurodegenerative diseases, such as “Pathways of neurodegeneration-multiple diseases”, “Amyotrophic lateral sclerosis” and “Huntington disease”. They were also enriched in several pathways related to pathogen infection, such as “Salmonella infection” and “Human T−cell leukemia virus 1 infection” (Figure 4D and Table S6).

**Figure 4.**
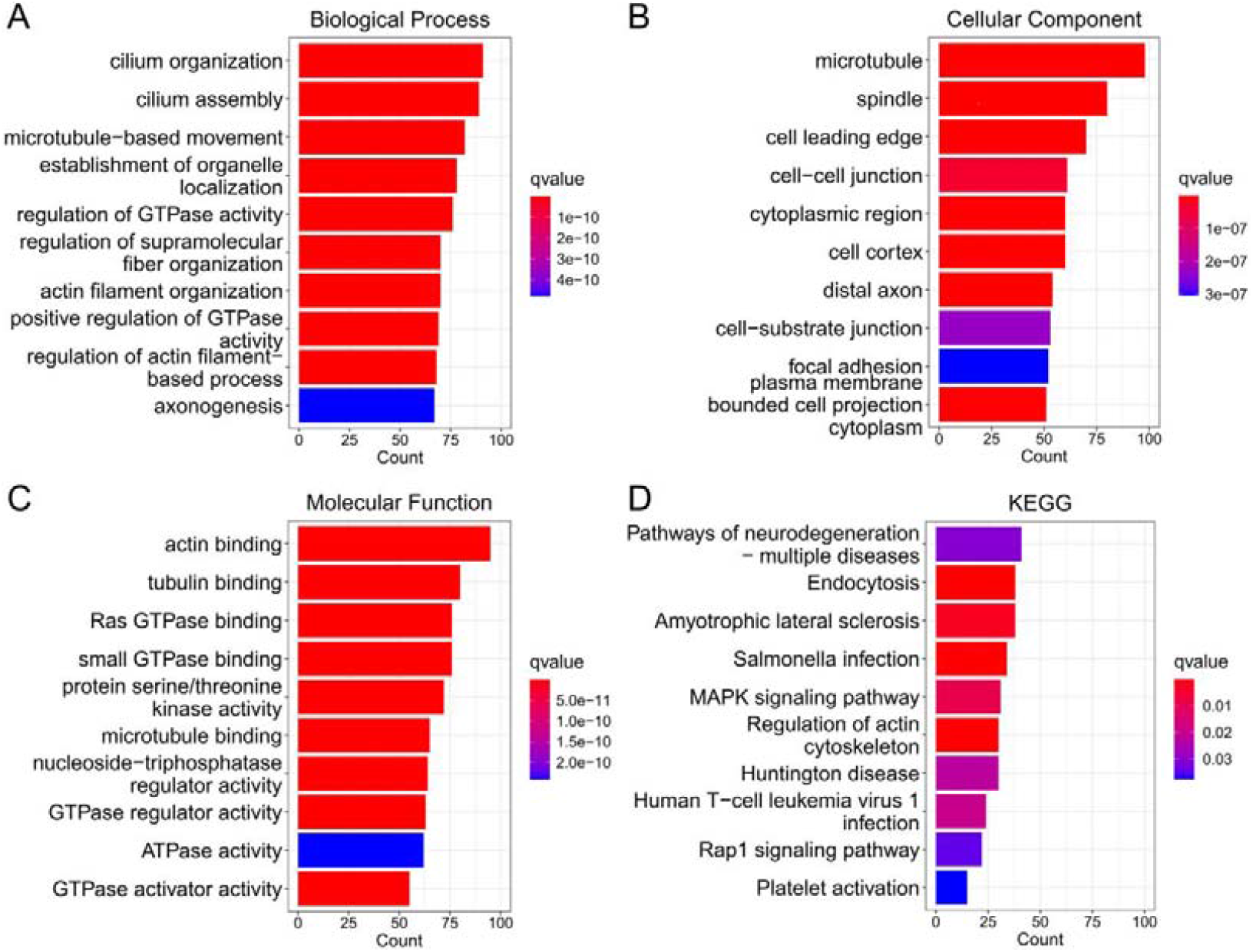
The functional enrichment analysis of the human proteins which were predicted to be cleaved by the coronavirus 3CL protease. Only top ten most enriched GO terms or KEGG pathways were shown. For more results, see Table S6. (A)-(C) refer to the GO enrichment analysis in the domain of Biological Process, Cellular Component, Molecular Function, respectively; (D) refers to the KEGG enrichment analysis.

### Construction of the online server 3CLP

To facilitate the usage of the RF model mentioned above, a user-friendly online server named 3CLP was built for predicting the cleavage sites of the coronavirus 3CL protease. The input of 3CLP is the protein sequences of either viral or host proteins in the fasta format; the prediction process would take several minutes depending on the number of protein sequences inputted; the outputs of 3CLP are the positions of the predicted cleavage sites, the SASA of the P1 and P1’, the motifs around the cleavage sites, and the score of the predicted cleavage sites which range from 0 to 1. The 3CLP is freely available to users without registration. The URL of 3CLP is http://www.computationalbiology.cn/3CLPHost/home.html.

## Discussion

This work systematically investigated the diversity of the cleavage sites of coronavirus 3CL protease on the polyprotein and found that the cleavage sites were highly conserved in multiple genera of the coronavirus. The AA preference at neighboring positions of the cleavage sites of the 3CL protease were similar to that reported in previous studies. For example, hydrophobic and small AAs were preferred at the P2 and P1’ position, respectively (Chuck et al., 2011). This preference enabled us to build the computational models of predicting the cleavage site based on the AA indexes instead of the AA identity.

The machine-learning-based methods and the homology-based methods have been developed to predict the cleavage sites of the coronavirus 3CL protease. Compared to the homology-based methods, the machine-learning-based methods can be used to predict the potential cleavage sites on host proteins, facilitating the studies of the virus-host interactions in viral infection. This study used the cleavage sites on polyproteins from 14 coronaviruses for modeling which were more than three times to that used in Kiemer’s study (Kiemer et al., 2004). Besides, this study used a very strict testing strategy by separating the dataset using the clustering method (Lu et al., 2021), which could reflect the ability of the model in predicting cleavage sites on polyproteins of novel viruses or on host proteins. In the independent testing on host proteins, although it only predicted one-third of the cleavage sites, the model achieved a high precision, suggesting its potential usage in predicting cleavage sites on host proteins.

Besides the cleavage on the viral polyproteins, the coronavirus 3CL protease can also cleave proteins involved in the host innate immune response such as NEMO and STAT2, thus evading the host immunity (Wang et al., 2016; Zhu et al., 2017a; Zhu et al., 2017b). For example, the 3CL protease of both the feline infectious peritonitis virus (FIPV) and porcine epidemic diarrhea virus (PEDV) can interrupt the type I interferon (IFN) signaling by cleaving the NEMO, which led to the reduction of type I IFN (Wang et al., 2016; Chen et al., 2019). This study predicted 1,420 potential cleavage sites in 1,079 human proteins. Besides the pathways related to pathogen infection, interestingly, these human proteins were also enriched in several KEGG pathways related to neurodegenerative diseases. Previous studies have shown that a large proportion of COVID-19 patients have developed the neurological symptoms and some patients even developed the Parkinsonism after the SARS-CoV-2 infection (Acharya et al., 2020; Cohen et al., 2020; Dewanjee et al., 2021; Fearon and Fasano, 2021; Taquet et al., 2021). Some patients infected by the MERS-CoV and SARS-CoV also presented severe neurological symptoms or complications (Lau et al., 2004; Tsai et al., 2004; Xu et al., 2005; Arabi et al., 2015; Kim et al., 2017). This study suggested that the neurological syndromes in patients infected by coronaviruses may be partly caused by the cleavage of critical proteins in nervous systems (such as actin and tubulin) by the viral 3CL protease. Further studies are needed to investigate the mechanism of neurological syndromes caused by the coronavirus.

There were some limitations in the study. Firstly, although the dataset was much larger than that used in previous studies, the dataset was still limited in size. The cleavage sites of the *Deltacoronavirus* were not included in the analysis. Nevertheless, the computational model developed here still showed high accuracy in both validations and testing, suggesting their potential usage in predicting the cleavage sites of the coronavirus 3CL protease. Secondly, the P1 position of the coronavirus 3C-like protease cleavage site was supposed to be highly conserved with Q, although there were a few cleavage sites with other residues in the P1 position. More experimental efforts are needed to determine the AA specificity of the coronavirus 3CL protease cleavage sites. Thirdly, only 12 experimentally-validated cleavage sites were used to test the model. More experimental validations are needed to evaluate the performance of the model.

## Conclusion

This work systematically investigated the diversity of the cleavage sites of the coronavirus 3CL protease, which help understand the specificity of the protease. A RF model and the related server 3CLP for predicting the cleavage sites of the coronavirus 3CL protease was built with high accuracy and predicted a total of 1,079 human proteins which may be cleaved by the coronavirus 3CL protease. The work not only provides an effective tool for identifying the cleavage sites of the protease, but also provides insights into the molecular mechanism underlying the pathogenicity of coronaviruses.

## Acknowledgements

This work was supported by the National Key Plan for Scientific Research and Development of China (2016YFD0500300) and Hunan Provincial Natural Science Foundation of China (2020JJ3006).

## References

Acharya A., Kevadiya B.D., Gendelman H.E., Byrareddy S.N. (2020) SARS-CoV-2 Infection Leads to Neurological Dysfunction. J Neuroimmune Pharmacol 15:167–173. doi: 10.1007/s11481-020-09924-9

Anand K., Ziebuhr J., Wadhwani P., Mesters J.R., Hilgenfeld R. (2003) Coronavirus main proteinase (3CLpro) structure: basis for design of anti-SARS drugs. Science 300:1763–1767.

Arabi Y.M., Harthi A., Hussein J., Bouchama A., Johani S., Hajeer A.H., et al. (2015) Severe neurologic syndrome associated with Middle East respiratory syndrome corona virus (MERS-CoV). Infection 43:495–501. doi: 10.1007/s15010-015-0720-y

Arya R., Kumari S., Pandey B., Mistry H., Bihani S.C., Das A., et al. (2021) Structural insights into SARS-CoV-2 proteins. J Mol Biol 433:166725. doi: 10.1016/j.jmb.2020.11.024

Blom N., Hansen J., Blaas D., Brunak S. (1996) Cleavage site analysis in picornaviral polyproteins: discovering cellular targets by neural networks. Protein Sci 5:2203–2216.

Chafekar A., Fielding B.C. (2018) MERS-CoV: Understanding the Latest Human Coronavirus Threat. Viruses 10. doi: 10.3390/v10020093

Chen B., Tian E.-K., He B., Tian L., Han R., Wang S., et al. (2020) Overview of lethal human coronaviruses. Signal Transduct Target Ther 5:89. doi: 10.1038/s41392-020-0190-2

Chen S., Tian J., Li Z., Kang H., Zhang J., Huang J., et al. (2019) Feline Infectious Peritonitis Virus Nsp5 Inhibits Type I Interferon Production by Cleaving NEMO at Multiple Sites. Viruses 12. doi: 10.3390/v12010043

Chuck C.-P., Chong L.-T., Chen C., Chow H.-F., Wan D.C.-C., Wong K.-B. (2010) Profiling of substrate specificity of SARS-CoV 3CL. PLoS One 5:e13197. doi: 10.1371/journal.pone.0013197

Chuck C.-P., Chow H.-F., Wan D.C.-C., Wong K.-B. (2011) Profiling of substrate specificities of 3C-like proteases from group 1, 2a, 2b, and 3 coronaviruses. PLoS One 6:e27228. doi: 10.1371/journal.pone.0027228

Cohen M.E., Eichel R., Steiner-Birmanns B., Janah A., Ioshpa M., Bar-Shalom R., et al. (2020) A case of probable Parkinson’s disease after SARS-CoV-2 infection. Lancet Neurol 19:804–805. doi: 10.1016/S1474-4422(20)30305-7

Crooks G.E., Hon G., Chandonia J.-M., Brenner S.E. (2004) WebLogo: a sequence logo generator. Genome Res 14:1188–1190.

Cui J., Li F., Shi Z.-L. (2019) Origin and evolution of pathogenic coronaviruses. Nat Rev Microbiol 17:181–192. doi: 10.1038/s41579-018-0118-9

Dewanjee S., Vallamkondu J., Kalra R.S., Puvvada N., Kandimalla R., Reddy P.H. (2021) Emerging COVID-19 Neurological Manifestations: Present Outlook and Potential Neurological Challenges in COVID-19 Pandemic. Mol Neurobiol. doi: 10.1007/s12035-021-02450-6

Fang S., Shen H., Wang J., Tay F.P.L., Liu D.X. (2010) Functional and genetic studies of the substrate specificity of coronavirus infectious bronchitis virus 3C-like proteinase. J Virol 84:7325–7336. doi: 10.1128/JVI.02490-09

Fearon C., Fasano A. (2021) Parkinson’s Disease and the COVID-19 Pandemic. J Parkinsons Dis 11:431–444. doi: 10.3233/JPD-202320

Fu L., Ye F., Feng Y., Yu F., Wang Q., Wu Y., et al. (2020) Both Boceprevir and GC376 efficaciously inhibit SARS-CoV-2 by targeting its main protease. Nat Commun 11:4417. doi: 10.1038/s41467-020-18233-x

Gralinski L.E., Bankhead A., Jeng S., Menachery V.D., Proll S., Belisle S.E., et al. (2013) Mechanisms of severe acute respiratory syndrome coronavirus-induced acute lung injury. mBio 4. doi: 10.1128/mBio.00271-13

Hartenian E., Nandakumar D., Lari A., Ly M., Tucker J.M., Glaunsinger B.A. (2020) The molecular virology of coronaviruses. J Biol Chem 295:12910–12934. doi: 10.1074/jbc.REV120.013930

Hu B., Guo H., Zhou P., Shi Z.-L. (2021) Characteristics of SARS-CoV-2 and COVID-19. Nat Rev Microbiol 19:141–154. doi: 10.1038/s41579-020-00459-7

Katoh K., Standley D.M. (2013) MAFFT multiple sequence alignment software version 7: improvements in performance and usability. Mol Biol Evol 30:772–780. doi: 10.1093/molbev/mst010

Kawashima S., Pokarowski P., Pokarowska M., Kolinski A., Katayama T., Kanehisa M. (2008) AAindex: amino acid index database, progress report 2008. Nucleic Acids Res 36:D202–D205.

Kiemer L., Lund O., Brunak S., Blom N. (2004) Coronavirus 3CLpro proteinase cleavage sites: possible relevance to SARS virus pathology. BMC Bioinformatics 5:72.

Kim J.E., Heo J.H., Kim H.O., Song S.H., Park S.S., Park T.H., et al. (2017) Neurological Complications during Treatment of Middle East Respiratory Syndrome. J Clin Neurol 13:227–233. doi: 10.3988/jcn.2017.13.3.227

Klemm T., Ebert G., Calleja D.J., Allison C.C., Richardson L.W., Bernardini J.P., et al. (2020) Mechanism and inhibition of the papain-like protease, PLpro, of SARS-CoV-2. EMBO J 39:e106275. doi: 10.15252/embj.2020106275

Larsen C.N., Sun G., Li X., Zaremba S., Zhao H., He S., et al. (2020) Mat_peptide: comprehensive annotation of mature peptides from polyproteins in five virus families. Bioinformatics 36:1627–1628. doi: 10.1093/bioinformatics/btz777

Lau K.-K., Yu W.-C., Chu C.-M., Lau S.-T., Sheng B., Yuen K.-Y. (2004) Possible central nervous system infection by SARS coronavirus. Emerg Infect Dis 10:342–344.

Lu C., Zhang Z., Cai Z., Zhu Z., Qiu Y., Wu A., et al. (2021) Prokaryotic virus host predictor: a Gaussian model for host prediction of prokaryotic viruses in metagenomics. BMC Biol 19:5. doi: 10.1186/s12915-020-00938-6

Mitternacht S. (2016) FreeSASA: An open source C library for solvent accessible surface area calculations. F1000Res 5:189. doi: 10.12688/f1000research.7931.1

Moustaqil M., Ollivier E., Chiu H.-P., Van Tol S., Rudolffi-Soto P., Stevens C., et al. (2021) SARS-CoV-2 proteases PLpro and 3CLpro cleave IRF3 and critical modulators of inflammatory pathways (NLRP12 and TAB1): implications for disease presentation across species. Emerg Microbes Infect 10:178–195. doi: 10.1080/22221751.2020.1870414

Pedregosa F., Varoquaux G., Gramfort A., Michel V., Thirion B., Grisel O., et al. (2011) Scikit-learn: Machine Learning in Python.

Satija N., Lal S.K. (2007) The molecular biology of SARS coronavirus. Ann N Y Acad Sci 1102:26–38.

Schechter I., Berger A. (1967) On the size of the active site in proteases. I. Papain. Biochem Biophys Res Commun 27:157–162.

Singh J., Litfin T., Paliwal K., Singh J., Hanumanthappa A.K., Zhou Y. (2021) SPOT-1D-Single: Improving the Single-Sequence-Based Prediction of Protein Secondary Structure, Backbone Angles, Solvent Accessibility and Half-Sphere Exposures using a Large Training Set and Ensembled Deep Learning. Bioinformatics. doi: 10.1093/bioinformatics/btab316

Singh O., Su E.C.-Y. (2016) Prediction of HIV-1 protease cleavage site using a combination of sequence, structural, and physicochemical features. BMC Bioinformatics 17:478. doi: 10.1186/s12859-016-1337-6

Snijder E.J., Decroly E., Ziebuhr J. (2016) The Nonstructural Proteins Directing Coronavirus RNA Synthesis and Processing. Adv Virus Res 96. doi: 10.1016/bs.aivir.2016.08.008

Stanley J.T., Gilchrist A.R., Stabell A.C., Allen M.A., Sawyer S.L., Dowell R.D. (2020) Two-stage ML Classifier for Identifying Host Protein Targets of the Dengue Protease. Pac Symp Biocomput 25:487–498.

Taquet M., Geddes J.R., Husain M., Luciano S., Harrison P.J. (2021) 6-month neurological and psychiatric outcomes in 236[379 survivors of COVID-19: a retrospective cohort study using electronic health records. Lancet Psychiatry 8:416–427. doi: 10.1016/S2215-0366(21)00084-5

Tsai L.-K., Hsieh S.-T., Chao C.-C., Chen Y.-C., Lin Y.-H., Chang S.-C., et al. (2004) Neuromuscular disorders in severe acute respiratory syndrome. Arch Neurol 61:1669–1673.

Tunyasuvunakool K., Adler J., Wu Z., Green T., Zielinski M., Žídek A., et al. (2021) Highly accurate protein structure prediction for the human proteome. Nature. doi: 10.1038/s41586-021-03828-1

Vuong W., Khan M.B., Fischer C., Arutyunova E., Lamer T., Shields J., et al. (2020) Feline coronavirus drug inhibits the main protease of SARS-CoV-2 and blocks virus replication. Nat Commun 11:4282. doi: 10.1038/s41467-020-18096-2

Wang D., Fang L., Shi Y., Zhang H., Gao L., Peng G., et al. (2016) Porcine Epidemic Diarrhea Virus 3C-Like Protease Regulates Its Interferon Antagonism by Cleaving NEMO. J Virol 90:2090–2101. doi: 10.1128/JVI.02514-15

WHO. (2021) WHO Coronavirus (COVID-19) Overview(https://covid19.who.int/).

Xu J., Zhong S., Liu J., Li L., Li Y., Wu X., et al. (2005) Detection of severe acute respiratory syndrome coronavirus in the brain: potential role of the chemokine mig in pathogenesis. Clin Infect Dis 41:1089–1096.

Yu G., Wang L.-G., Han Y., He Q.-Y. (2012) clusterProfiler: an R package for comparing biological themes among gene clusters. OMICS 16:284–287. doi: 10.1089/omi.2011.0118

Zhu X., Chen J., Tian L., Zhou Y., Xu S., Long S., et al. (2020) Porcine Deltacoronavirus nsp5 Cleaves DCP1A To Decrease Its Antiviral Activity. J Virol 94. doi: 10.1128/JVI.02162-19

Zhu X., Fang L., Wang D., Yang Y., Chen J., Ye X., et al. (2017a) Porcine deltacoronavirus nsp5 inhibits interferon-β production through the cleavage of NEMO. Virology 502:33–38. doi: 10.1016/j.virol.2016.12.005

Zhu X., Wang D., Zhou J., Pan T., Chen J., Yang Y., et al. (2017b) Porcine Deltacoronavirus nsp5 Antagonizes Type I Interferon Signaling by Cleaving STAT2. J Virol 91. doi: 10.1128/JVI.00003-17

